# Non-muscle myosin 2 can incorporate into established filaments in cells without an assembly competence domain

**DOI:** 10.1101/2024.07.07.602405

**Authors:** Kehan Wu, Hiral Patel, Huini Wu, Melissa A. Quintanilla, Margaret A. Bennett, Stefano Sala, Jordan R. Beach

## Abstract

Myosin 2 dynamically assembles into filaments that exert force on the actin cytoskeleton. To form filaments, myosin 2 monomers transition between folded and unfolded states. Monomer unfolding exposes an extended coiled-coil that interacts with other monomers in parallel and antiparallel fashions, enabling bipolar filament formation. A C-terminal domain of the coiled-coil, termed assembly competence domain (ACD), has been repeatedly identified as necessary for filament assembly. Here, we revisit ACD contribution when full-length filaments are present. Non-muscle myosin 2A lacking the ACD (ΔACD) initially appears diffuse, but triton extraction of cytosolic fraction reveals cytoskeletal association. Disruption of the folded monomer enhances the cytoskeletal fraction, while inhibition of endogenous filament assembly appears to reduce it. Finally, high resolution imaging of endogenous and exogenous bipolar filamentous structures reveals highly coincident signal, suggesting ΔACD constructs co-assemble with endogenous myosin 2A filaments. Our data demonstrate that while the ACD is required for de novo filament assembly, it is not required for monomers to recognize and associate with established filaments in cells. More broadly, this highlights the existence of distinct mechanisms governing myosin 2 monomer assembly into nascent filaments, and monomer recognition and association with established filaments to maintain steady-state contractile networks.

## Introduction

Myosin 2 motor proteins are the dominant contractile motor proteins in mammalian cells. To function, myosin 2 monomers assemble into bipolar filaments that engage filamentous actin to drive contraction (reviewed in (1, 2).) Defining the process of myosin 2 filament formation informs how cells build and maintain contractile network dynamics to drive an array of cellular processes, including migration and division. In non-muscle cells, roughly one-to two-thirds of the myosin 2 exists in the filamentous state (3–5). Although filaments are continually being assembled and disassembled throughout the cell, it is likely that many filaments exist for extended periods with a steady-state exchange of monomers moving into and out of the filament. Therefore, while identifying mechanisms by which myosin 2 monomers assemble nascent filaments is critical for understanding how contractile networks are built, identifying mechanisms by which myosin 2 monomers recognize and associate with established filaments is critical for understanding how contractile networks are maintained.

Myosin 2 monomers are hexameric ensembles of three components: two myosin heavy chains (MHC), two regulatory light chains (RLC), and two essential light chains (ELC). MHCs consist of a motor domain, a neck region with two light-chain binding IQ motifs, an extended alpha-helix that dimerizes into an extended coiled-coil “tail”, and terminate in a short non-helical tailpiece (Fig. 1A) (1). The coiled-coil is imperative for filament assembly and a focus of this work.

**Fig. 1.**
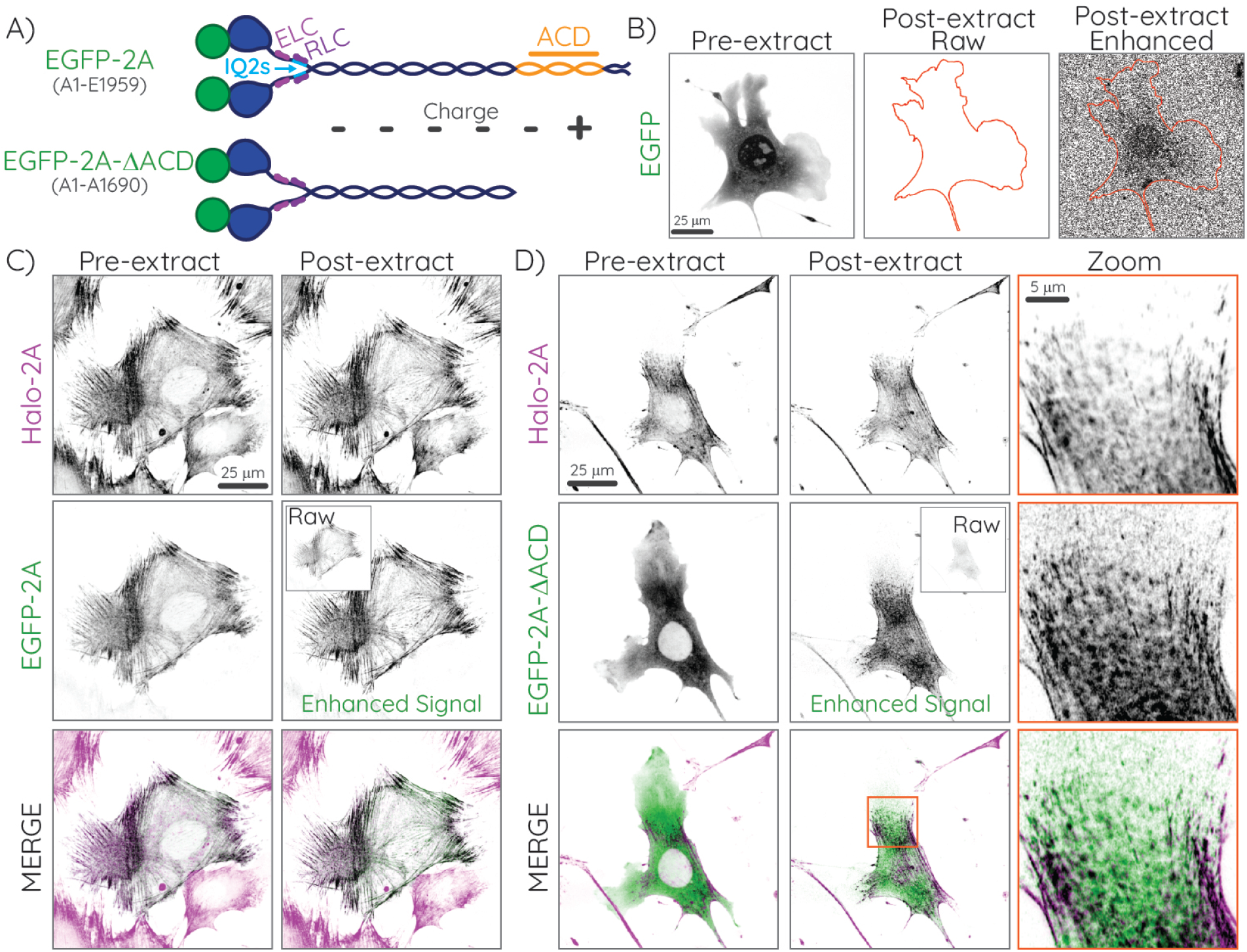
EGFP-2A-ΔACD colocalizes with endogenous myosin 2A networks upon extraction of the cytosolic fraction. A) Cartoon of full-length EGFP-2A (top) and EGFP-2A-ΔACD (bottom) with amino acids in parentheses. The ACD is displayed in orange. Minus and plus signs indicate the net charged regions along the coiled-coil tail. B - D) Summed confocal z-projections. “Raw” panels and insets are scaled for intensity identical to pre-extraction images, while “Enhanced” images are scaled independently. Fibroblasts transiently expressing diffuse EGFP imaged pre- and post-extraction. C and D) Halo-2A fibroblasts expressing EGFP-2A (C) or EGFP-2A-ΔACD (D) imaged pre- and post-extraction. The rightmost column in D displays zoomed insets of the orange box in the EGFP-2A-ΔACD post-extraction merged image. See also Movie S1.

To prevent spurious filament assembly, mammalian myosin 2 monomers can be sequestered into a folded, inactive state termed the “10S” (6). In the 10S state, the motor domains autoinhibit by docking on one another and fold back to bind along the N-terminus of the coiled-coil, creating the interacting heads motif (IHM) (7, 8). The coiled-coil tail also folds twice to wrap around the IHM, further stabilizing the sequestered state (9–11). The presence of unphosphorylated RLCs is critical for 10S formation and stability (6, 12). Phosphorylation of the RLC at T18/S19, or deletion of the second IQ motif (ΔIQ2) where the RLC binds, destabilizes the 10S and permits unfolding into the assembly-competent “6S” monomer (13, 14). This unfolding exposes the coiled-coil tail. Surface-exposed residues along the coiled-coil display alternating negative and neutral\ charge regions, with a positive charge region near the C-terminus (Fig. 1A) (15–18). These alternating charge regions enable staggered electrostatic interactions that stabilize parallel and antiparallel monomer interactions to promote bipolar filament assembly (18, 19). Therefore, the standard assembly model is that the folded, inactive 10S diffuses throughout the cytoplasm until it is phosphorylated by RLC kinases to induce unfolding into the assembly-competent activated 6S that subsequently assembles into filaments. Alternative non-mutually exclusive assembly models have also been proposed, which suggest that the 10S monomer or 10S dimers first interact with established filaments, and then unfold into the 6S while already filament-associated (20).

Foundational studies to establish mechanisms of assembly have been performed using purified proteins. While examining both full-length and tail fragments of various myosin 2s, these studies defined an assembly competence domain (ACD) in 1 the C-terminus of the coiled-coil, which universally includes the lone positive charge region (21–25). Truncation studies deleting the ACD (ΔACD) or consisting solely of the ACD established that the ACD is both necessary and sufficient for paracrystal formation or insolubility, suggestive of it being necessary and sufficient for filament assembly. However, these studies predominantly examined homogenous populations for their ability to assemble, and did not examine mixed populations of full-length myosin 2 with truncated myosin 2. Therefore, these studies were examining capacity for monomers to assemble new filaments, not the capacity for monomers to recognize and associate with established filaments in steady-state contractile networks.

Parallel to the in vitro work, several studies examined tail truncations in *Dictyostelium* and mammalian cells (3, 23, 26–29).

Over-expression of EGFP-tagged myosin 2 with C-terminal truncations that removed all or part of the ACD was reported to abolish assembly and result in diffuse cytosolic myosin 2. This suggested that even in the presence of endogenous full-length myosin 2, the ACD is necessary for recognition and association with established filaments. Reciprocal to ΔACD constructs preventing assembly, ΔIQ2 constructs that prevent RLC binding are thought to be constitutively filamentous (3, 30). This is due to the inability to form the autoinhibited 10S monomer, thus locking the molecule into the unfolded 6S which readily assembles into filaments. A double deletion ΔIQ2ΔACD construct also appeared cytosolic when expressed in mammalian cells, but displayed slower diffusion kinetics than the ΔACD alone, consistent with it being locked into the unfolded 6S monomer but unable to assemble (3).

Here, we revisit myosin 2 truncation and deletion mutants using the mammalian non-muscle myosin 2A isoform (hereafter myosin 2A) to test models of filament assembly and maintenance in cells. Contrary to previous studies, we find that ACD deletion does not prevent recognition and association with established full-length myosin 2 filaments in cells. Moreover, double deletion of both the ACD and the IQ2 domains enhances filament association. Inhibition of endogenous filament assembly with either ROCK inhibition or genetic ablation of myosin 2A (*Myh9*) demonstrates this filament association is dependent on the presence of endogenous filaments. Finally, high resolution imaging reveals highly coincident exogenous and endogenous bipolar filamentous structures, consistent with these deletion constructs co-assembling into full-length endogenous myosin 2 filaments.

## Results

### Retention of myosin 2A filament association upon ACD deletion

To test the contribution of the ACD to the steady-state incorporation of myosin 2A monomers into established filaments, we expressed EGFP, EGFP-myosin 2A (“EGFP-2A”), or EGFP-myosin 2A-ΔACD (“EGFP-2A-ΔACD”) in a mouse fibroblast cell line (JR20s) where endogenous myosin 2A is labeled with a HaloTag (Halo-2A; Fig. 1A and Fig. S1). While the EGFP signal appeared diffuse (Fig. 1B; Pre-extract), myosin 2A displayed cytoskeletal localization that was highly coincident with the endogenous Halo-myosin 2A signal (Fig. 1C; Pre-extract). Similar to EGFP, and in agreement with previous publications, EGFP-2A-ΔACD displayed a diffuse signal in fibroblasts, with no discernible cytoskeletal appearance (Fig. 1B, 1D; Pre-extract). However, extraction of the cells with a physiological buffer containing Triton X-100, which releases the cytosolic EGFP (Fig. 1B; Post-extract) and cytosolic myosin 2 fractions but retains filamentous myosin 2, revealed a cytoskeletal myosin 2-like distribution of EGFP-2A-ΔACD (Fig. 1D; Post-extract; Movie S1). While the post-extracted raw EGFP-2A-ΔACD signal intensity was reduced relative to EGFP-2A (Fig. 1; “Raw” insets), the enhanced signal was highly coincident with the endogenous Halo-2A signal (Fig. 1B - 1D; post-extraction zoomed insets). These data suggest that while the majority of the EGFP-2A-ΔACD molecules are monomeric, a fraction might associate with and incorporate into endogenous myosin 2A filaments.

### Inhibition of 10S monomer increases assembly of EGFP-2A-ΔACD

We hypothesized that EGFP-2A-ΔACD follows a canonical assembly process, whereby RLC phosphorylation inhibits the folded 10S monomer, generating unfolded 6S monomers that transiently assemble into established endogenous myosin 2 filaments. To explore this, we used a previously published construct that inhibits 10S formation by deleting the RLC-binding IQ2 motif, termed EGFP-2A-ΔIQ2ΔACD (Fig. 2A)(3). Previous studies theorized that this construct would reside in the 6S intermediate, unable to form the folded 10S and unable to assemble into filaments. When expressed in our fibroblasts, EGFP-2A-ΔIQ2ΔACD displayed a diffuse cytosolic signal pre-extraction in a subpopulation of the cells (Fig. 2B; Pre-extract). However, many cells also displayed clear cytoskeletal structures prior to any extraction (Fig. 3C and Movies S1 and S2). Upon triton-extraction, we readily observed a robust cytoskeletal, myosin 2-like network in all cells (Fig. 2B; Post-extract). Similar residual cytoskeletal signal upon triton extraction was observed for both EGFP-2A-ΔACD and -ΔIQ2ΔACD in cell types previously used to investigate these deletion/truncation constructs (Hela and U2OS cells; Fig. S2), demonstrating our observations are not cell-type dependent. These data indicate that deletion of both the ACD and IQ2 motif does not prevent the incorporation of EGFP-2A into endogenous filaments. To quantify these observations, we developed an analysis pipeline to measure the intracellular EGFP intensity pre- and post-extraction based on a cell mask created using the consistent endogenous Halo-2A signal (Fig. 3A and Fig. S3). The post-extraction to pre-extraction ratio serves as a proxy for filamentous protein (i.e. insoluble fraction). As expected, the insoluble fraction for the EGFP control was near zero, consistent with its cytosolic localization (Fig. 3B). In agreement with previous reports (3–5), approximately half of the EGFP-2A molecules were in the filamentous form (Fig. 3B). While about 10 percent of EGFP-2A-ΔACD was incorporated into filaments, the insoluble levels of EGFP-2A-ΔIQ2ΔACD were comparable to those of full-length EGFP-2A. Plotting the pre-extraction EGFP intensities versus the insoluble fraction for each construct revealed that their insoluble fractions were independent of their expression levels (Fig. S1B). These data quantitatively confirm that the ACD is not requisite for cytoskeletal association, and that inhibition of 10S formation via IQ2 deletion increases myosin 2A filament assembly even in the absence of an ACD.

**Fig. 2.**
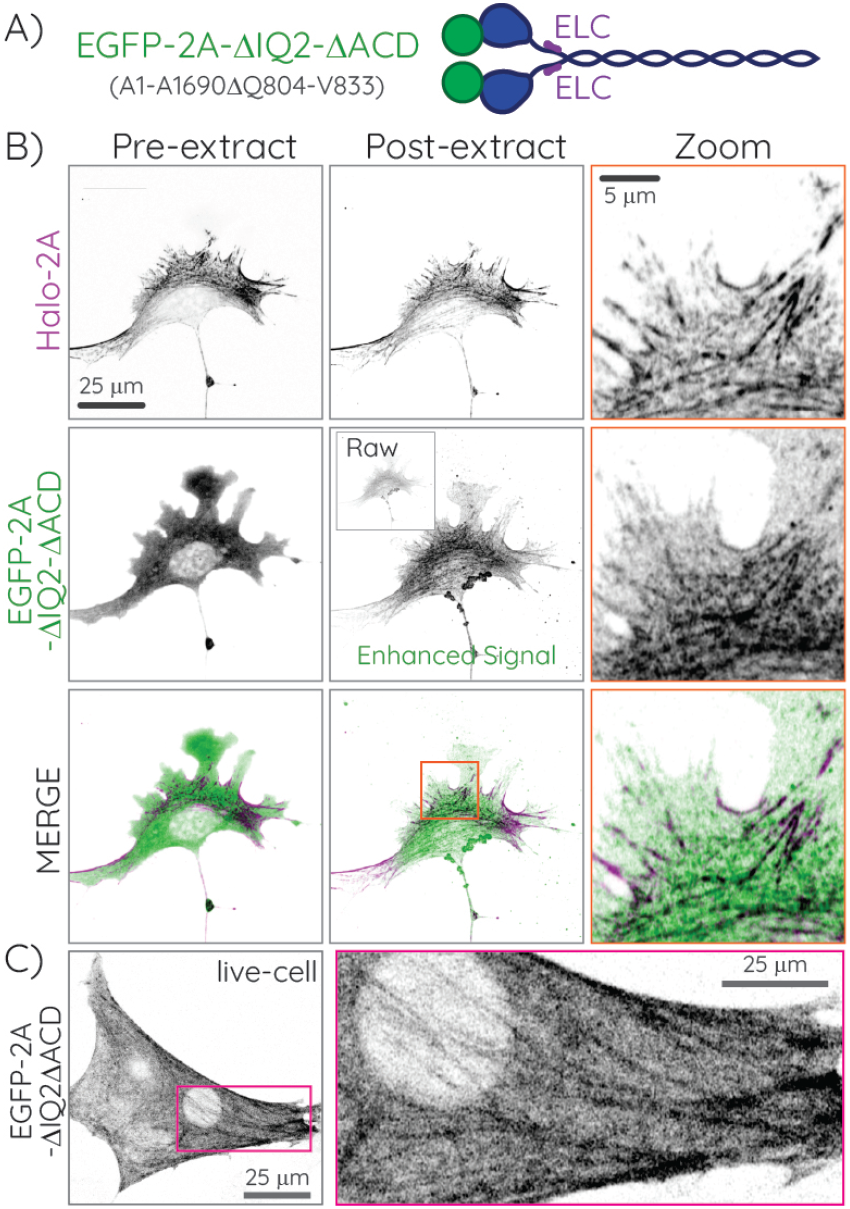
Deletion of both the IQ2 motif and ACD enhances association with endogenous myosin filaments. A) Cartoon of EGFP-2A-ΔIQ2ΔACD with amino acids in parentheses. Note the loss of RLC binding via IQ2 deletion. B) Summed confocal z-projection of Halo-2A fibroblasts transiently expressing EGFP-2A-ΔIQ2ΔACD. “Raw” inset is scaled for intensity identical to pre-extraction image, while “Enhanced” image is scaled independently. The rightmost column displays zoomed insets of the orange box in EGFP-2A-ΔIQ2ΔACD post-extraction merged image. C) Live cell imaging of the EGFP-2A-ΔIQ2ΔACD construct in fibroblasts. The zoomed inset (right) of the magenta box on the left shows the cytoskeletal localization of EGFP-2A-ΔIQ2ΔACD. See also Movies S1 and S2.

**Fig. 3.**
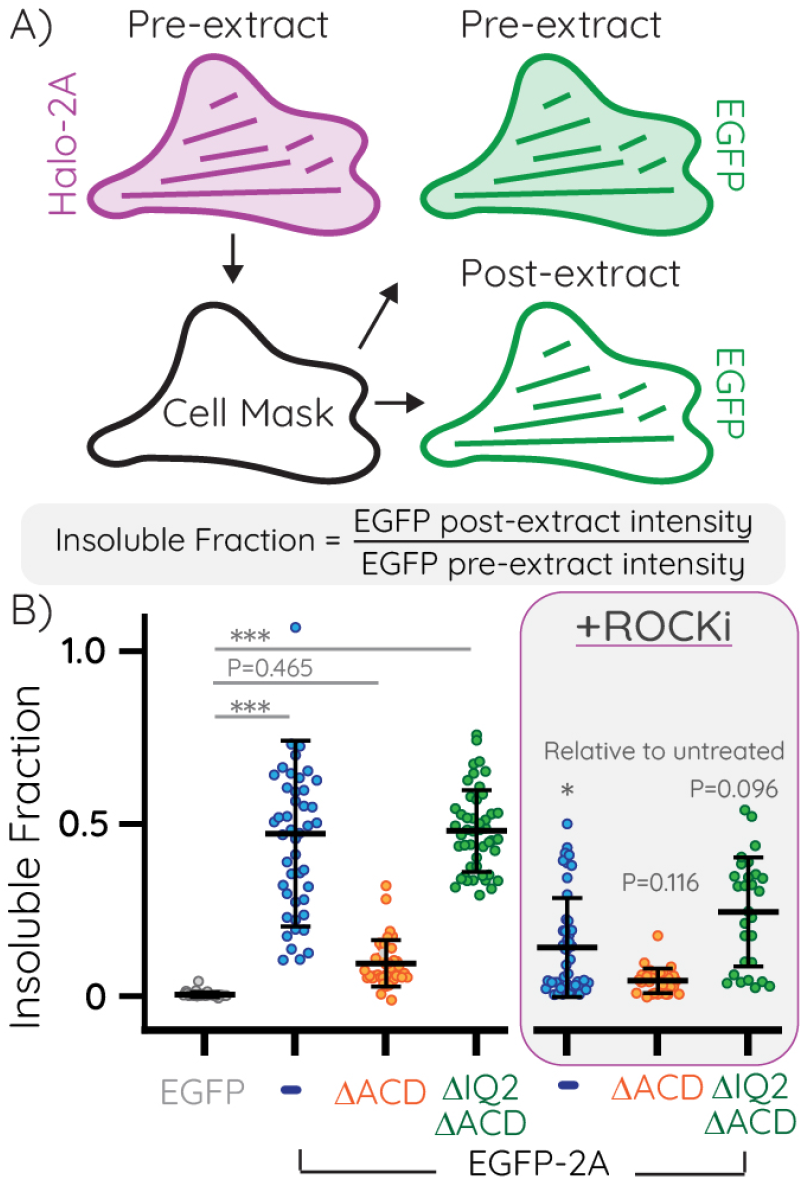
Quantification of myosin 2A insoluble fractions in triton fractionation assays. A) Cartoon of the analysis pipeline (for details, see Supplementary Fig. S3). Images of Halo-2A fibroblasts transiently expressing EGFP-2A constructs pre- and post-extraction were used for analysis. The consistent endogenous Halo-2A signal was used to generate a cell mask, which was applied to the pre- and post-extraction EGFP images. The insoluble fraction, as a correlate for filament assembly, was calculated as the ratio of post-extraction intensity to pre-extraction intensity. B) Comparison of the insoluble fractions for EGFP or EGFP-2A constructs without (left) or with (right) ROCK inhibition using 10 µM Y27632 for 30 minutes. Data points indicate individual cells with error bars indicating mean +/-SD. Statistical comparisons were made using nested one-way ANOVA and Tukey’s multiple comparison tests. P values are indicated for “not significantly different” comparisons between untreated and ROCK inhibited data sets. See Table S1 for replicates and full statistical comparisons. Data points outside the scale were removed from the graph but included in statistical analyses.

### Filament association of EGFP-2A-ΔACD and -ΔIQ2ΔACD is dependent on endogenous myosin 2A filaments

To test our hypothesis that the EGFP-2A-ΔACD and EGFP-2A-ΔIQ2ΔACD truncation constructs co-assemble into endogenous filaments, we used two parallel assays to reduce endogenous myosin filament levels. First, we inhibited the dominant RLC kinase in fibroblasts, rho-associated coiled-coil containing kinase (ROCK), using the small molecule Y27632. As expected, mild ROCK inhibition reduced EGFP-2A assembly by about four-fold, and reduced both EGFP-2A-ΔACD and EGFP-2A-ΔIQ2ΔACD assembly by about two-fold (Fig. 2B). These data are consistent with the majority of the insoluble fraction for these constructs co-assembling with the endogenous myosin 2A.

Our second approach to reducing endogenous filament levels was to genetically ablate endogenous myosin 2A (*Myh9* gene) in fibroblasts using CRISPR/Cas9. These fibroblasts predominantly express myosin 2A, with little expression of the myosin 2B and myosin 2C isoforms. Therefore, despite the ability of these isoforms to co-assemble (31, 32), deletion of myosin 2A should remove the majority of endogenous myosin 2, significantly reducing the filamentous appearance of EGFP-2A-ΔACD and -ΔIQ2ΔACD. Transient expression and triton extraction of full-length EGFP-2A in myosin 2A-KO cells revealed that about fifty percent was filamentous (Fig. 4A-B), similar to what we observed in wild-type fibroblasts (Fig. 3B). For EGFP-2A-ΔACD, we again observed a diffuse signal pre-extraction and little residual signal post-extraction (Fig.4A-B). For the EGFP-2A-ΔIQ2ΔACD construct, filament incorporation was more apparent than EGFP-2A-ΔACD, with a higher insoluble fraction post-extraction compared to EGFP-2A-ΔACD (Fig. 4A-B). However, this was reduced about two-fold relative to the insoluble fraction of EGFP-2A-ΔIQ2ΔACD in wild-type fibroblasts (Fig. 3B). Notably, EGFP-2A-ΔIQ2ΔACD may be slightly less impacted by ROCK inhibition relative to EGFP-2A, as it does not have any RLC binding sites and therefore is not a direct target of ROCK. Our data show that while EGFP-2A-ΔACD and EGFP-2A-ΔIQ2ΔACD still assemble into filaments, their ability to do so is significantly reduced in the absence of endogenous myosin filaments.

**Fig. 4.**
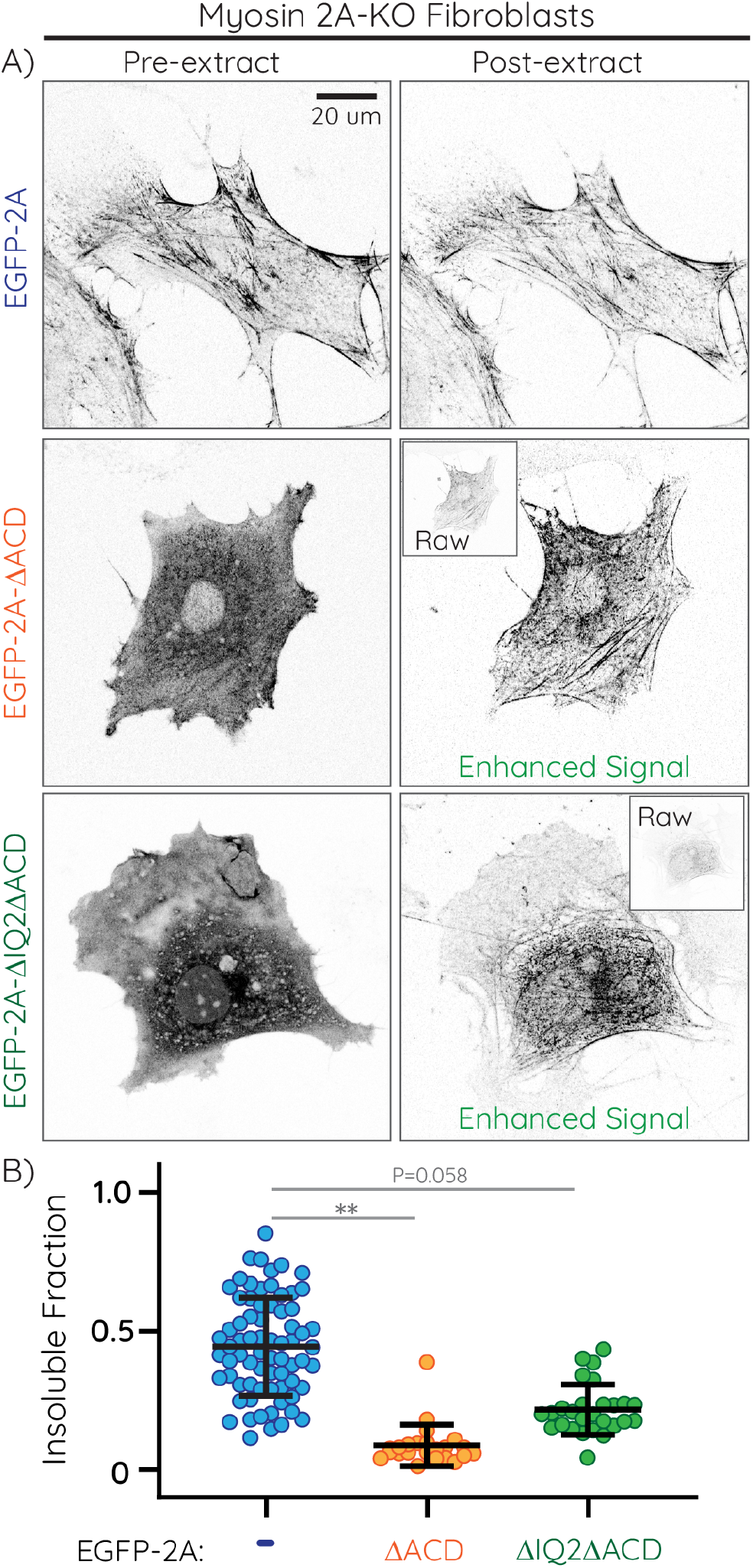
Myosin 2A KO reduces the assembly of EGFP-2A-ΔIQ2ΔACD. A) Summed confocal z-projection of myosin 2A KO fibroblasts transiently expressing EGFP-2A (top) EGFP-2A-ΔACD (middle) and EGFP-2A-ΔIQ2ΔACD (bottom) pre and post extraction. The EGFP signal was used to generate a cell mask, which was applied to the pre- and post-extraction images. “Raw” inset is scaled for intensity identical to pre-extraction image, while “Enhanced” image is scaled independently. B) Comparison of the insoluble fractions for EGFP-2A constructs. Data points indicate individual cells with error bars indicating mean +/-SD. Statistical comparisons were made using nested one-way ANOVA and Tukey’s multiple comparison tests. See Table S1 for replicates and full statistical comparisons.

### High resolution imaging of endogenous and exogenous myosin 2A is consistent with a co-assembly model

Finally, we performed high resolution imaging to investigate the co-assembly of endogenous and exogenous myosin 2A. Specifically, we used Zeiss Airyscan imaging with joint deconvolution processing, providing a sub 100 nm theoretical lateral resolution, sufficient to observe individual myosin 2 filaments or small filament stacks with fluorophore-tagged N-termini creating puncta approximately 300 nm apart (Fig. 5A). Imaging of extracted Halo-2A fibroblasts that express either EGFP-2A, EGFP-2A-ΔACD or EGFP-2A-ΔIQ2ΔACD demonstrated highly coincident bipolar structures consisting of endogenous Halo-2A and exogenous EGFP-2A constructs (Fig. 5B - 5D). These results are consistent with a co-assembly model. Collectively, these data argue that removal of the ACD does not preclude monomer recognition and association with established myosin 2 filaments, and that disruption of the 10S increases filament assembly even in the absence of an ACD.

**Fig. 5.**
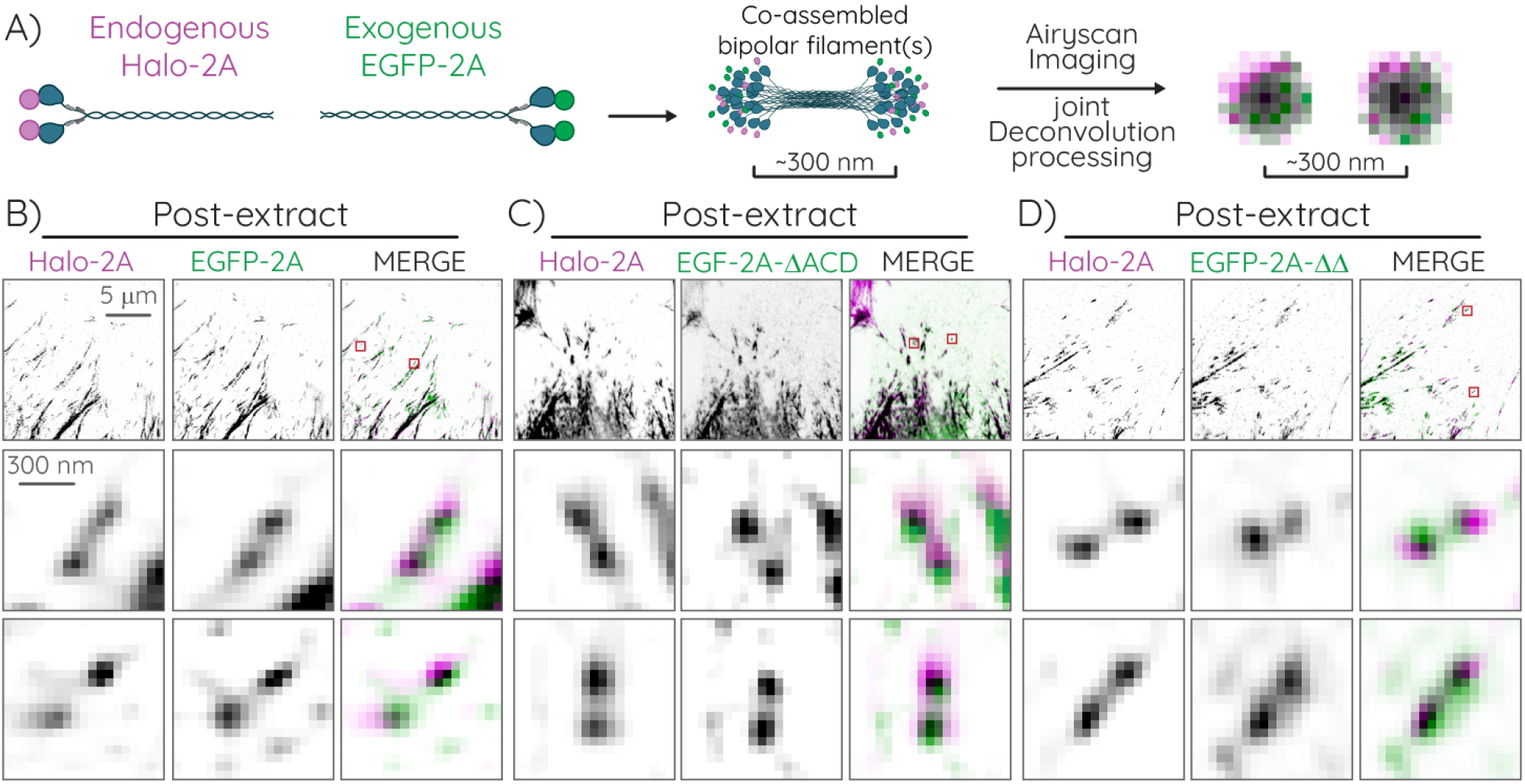
High-resolution imaging reveals likely co-assembled endogenous and exogenous myosin 2A filaments. A) Cartoon of co-assembled endogenous and exogenous myosin 2A filaments imaged with at high spatial resolution. Halo-2A (magenta) monomers co-assembled into bipolar filaments with EGFP-2A monomers (green) should appear as two puncta approximately 300 nm apart, with significant magenta and green overlap, when imaged with an Airyscan and processed with “joint Deconvolution” (see Methods). B-D) Post-extraction images of Halo-2A fibroblasts expressing the indicated EGFP-2A construct. Top row displays individual and merged myosin 2A channels in the lamellar region of the cell. The bottom two rows show the zoomed insets of the red boxes in the top row of Halo-2A bipolar structures, along with the corresponding myosin 2A signal, which display significant overlap, consistent with co-assembly.

## Discussion

Here, we provide novel insight into how myosin 2 monomers recognize and incorporate into established myosin 2 filaments. We posit that there are potentially unique mechanistic differences between monomers assembling into a nascent filament (Fig. 6; top) and monomers recognizing and incorporating into established filaments (Fig. 6; bottom). This necessitates unique, but not mutually exclusive, molecular models to test. To assemble into nascent filaments, we maintain that the ACD is absolutely requisite. Definitive experiments with purified protein and modeling inform us that the positively charged regions within the ACD are needed to stabilize electrostatic interactions between myosin 2 monomers in both a parallel and anti-parallel manner (21–25). These interactions could occur between 10S dimers that subsequently unfold into nascent filaments (20) or occur between unfolded 6S monomers that encounter each other in the cytosol. Regardless, the positively charged region within the ACD, and therefore the ACD itself, is required for de novo assembly.

**Fig. 6.**
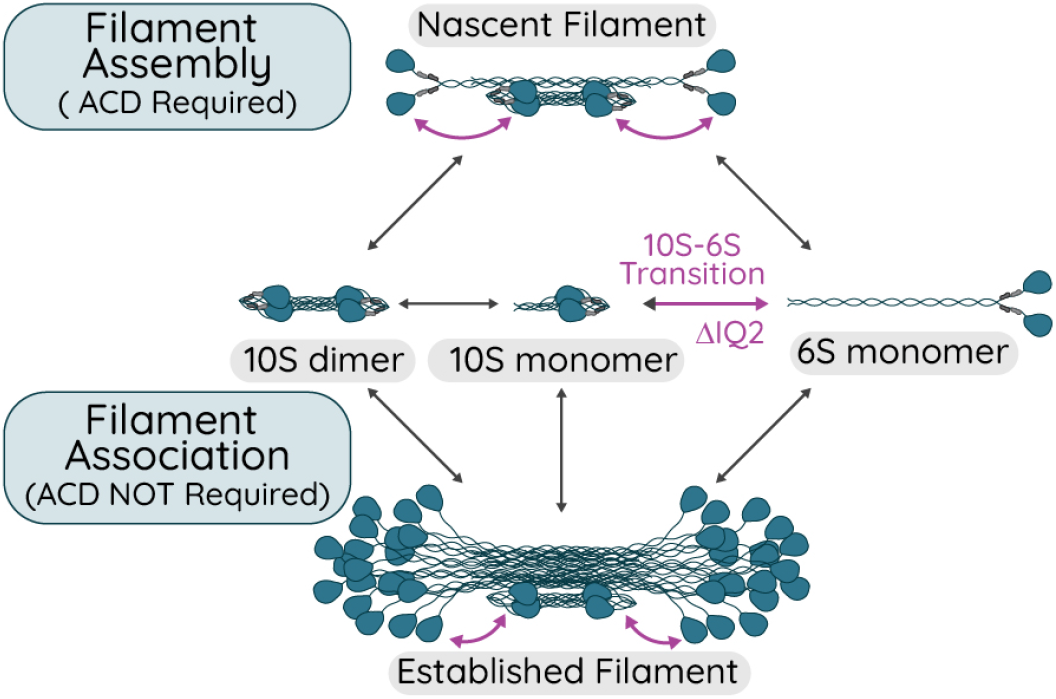
Model of Myosin 2 Filament Assembly and Association. See Discussion.

In contrast, we argue that the ability of myosin monomers to recognize and incorporate into an established filament is a distinct concept, which has often either been overlooked or incorrectly grouped into the former concept of nascent filament assembly. While there is likely significant overlap in the molecular interactions involved in filament assembly and filament association, there could also be exclusive interactions that enable association but are not sufficient for assembly. By removing the ACD in our study, we were able to distinguish between filament assembly and filament recognition. We demonstrate that even in the absence of an ACD, myosin 2 molecules can associate with and incorporate into established filaments composed of full-length myosin 2. Furthermore, through additional removal of the IQ2 motif and inhibition of the 10S, we demonstrate that the extended 6S monomer, even without an ACD, can readily incorporate into established filaments composed of full-length myosin 2. We note that our results do not invalidate non-mutually-exclusive models suggesting 10S monomers or 10S dimers can associate with established filaments and then unfold and incorporate into filaments.

Our work elicits consideration of the scale of de novo assembly of new myosin 2 filaments relative to the maintenance of established myosin 2 filaments. Certainly, de novo assembly of myosin 2 filaments is critical throughout cell biology. However, the scale of de novo assembly contribution may be skewed towards processes involving significant morphological changes where many new filaments need to be generated (e.g. cytokinesis, muscle hypertrophy, etc.). In contrast, steady-state maintenance of established filaments is likely a dominant pathway in many “stationary” cells throughout the body. We do not know what the lifetime of an established filament is. However, we do know that the half-times of recovery for non-muscle myosin 2 in FRAP studies are about 0.5 - 1.5 minutes, indicating that monomers are readily exchanging into and out of established filaments (3, 33). Surprisingly, rapid exchange kinetics have also recently been reported for cardiac myosin 2 (34). Therefore, in steady-state contractile environments (e.g., stress fibers or sarcomeres), it is possible that the original monomers in the filament are entirely replaced multiple times during the filament’s lifetime. One could even ponder, a la the Ship of Theseus, “if all monomers of a filament are replaced is it still the same filament?” Regardless of the philosophical answer, the collective evidence argues that replacing monomers within filaments happens and is critical to maintaining physiological levels of contractility.

Currently, we can only speculate on the precise molecular interactions enabling our truncation/deletion constructs to associate with established contractile networks. We hypothesize that the negatively charged regions in the proximal tail of the EGFP-2A-ΔACD and -ΔIQ2ΔACD constructs are able to recognize and interact with the positively charged regions in the ACD of the full-length myosin 2 already present in established filaments. This suggests that these positively charged regions are dynamically and transiently available for monomer recruitment into established filaments and not permanently engaged in electrostatic interactions while in the filament. Alternatively, or additionally, low-affinity interactions that stabilize the folded 10S (11) could facilitate hetero-molecular interactions between monomeric and filamentous myosin 2, thereby recruiting the monomer.

In addition to interacting with endogenous myosin filaments, the retention of EGFP-2A-ΔIQ2ΔACD after triton extraction may be due to additional interactions. The most likely interaction is with actin, as the absence of RLCs should disrupt the IHM, enabling actin binding. Our data might even shed light on this possibility: First, we often see higher background intensity post-extraction with EGFP-2A-ΔIQ2ΔACD (Fig. 2B and Fig. 4A). This could be due to a low actin-binding capacity by these monomers. Second, if EGFP-2A-ΔIQ2ΔACD insolubility was entirely dependent on endogenous filament assembly, ROCK inhibition would have impacted both EGFP-2A and EGFP-2A-ΔIQ2ΔACD similarly. However, EGFP-2A insolubility decreased four-fold while EGFP-2A-ΔIQ2ΔACD decreased only two-fold. Some of this resistance to ROCK inhibition could be due EGFP-2A-ΔIQ2ΔACD not being a direct target of ROCK, leaving unfolded molecules that can bind actin even with limited endogenous filaments present.

It is worth noting that several previous studies, which similarly involved over-expressing GFP-tagged myosin 2 truncations in mammalian cells, did not observe filamentous structures (3, 35). Minor sequence differences in the creation of the truncated ΔACD construct might account for some discrepancy, but we think this is unlikely. We performed our triton extraction assay in the same cell types used in those earlier studies (e.g. HeLa and U2OS) and observed similar cytoskeletal localization of both EGFP-2A-ΔACD and -ΔIQ2ΔACD, ruling out a cell-type specific observation. Considering our quantitative analysis suggests that over 90% of EGFP-2A-ΔACD is cytosolic, we believe that this cytosolic component masked the remaining filament-incorporated component in earlier studies, rendering it undetectable without removal of the cytosolic fraction. This argues that any fluorophore-tagged protein that initially appears diffuse should be interpreted carefully, and researchers should consider experiments to remove cytosolic fractions to determine if underlying stable associations are present.

From a technical perspective, we highlight that myosin 2-ΔACD should not be used as a negative control for assembly assays where full-length myosin 2 and/or actin are present. Even in the absence of full-length myosin 2, myosin 2-ΔACD constructs could bind actin and present as filamentous. Further, while there may be differences in the assembly mechanisms between non-muscle myosin 2 and smooth/striated myosin 2 paralogs, it is likely that our observations apply to all myosin 2 paralogs. Therefore, experimentation with any ΔACD construct should be performed with caution.

In conclusion, our key observation that the ACD is not a requisite domain for recognition and incorporation into established filaments highlights that the terms “filament assembly” and “filament association” are not synonymous terms. We need to consider potentially exclusive mechanisms by which monomers recognize and incorporate into established filaments that are unique from mechanisms by which monomers are able to assemble de novo filaments. Isolating these concepts is experimentally challenging. However, at minimum, we argue that the field should be careful not to conflate these concepts when interpreting and discussing data.

## Materials and Methods

### Cell culture and transfection

Halo-2A (“H2A2”), myosin 2A KO, U2OS and Hela cells were cultured in DMEM (10-013-CV, Corning) supplemented with 10% FB Essence (10803-034, Avantor Seradigm) and 1% antibiotic-antimycotic (30-004-Cl, Corning) at 37°C and 5% CO2. To generate CRISPR knock-in lines, 1 million cells were transfected with 5 µg each of the target and donor plasmid using the LipoD293 (SL100668, SignaGen) system. Twenty four hours before triton experiments, 400,000 cells (H2A2, U2OS or Hela) were transfected with 3 µg (GFP) or 4 µg (EGFP-2A constructs) plasmid DNA using a Neon electroporation system (ThermoFischer Scientific) with a single 20 ms pulse of 1600V and plated on glass coverslips. Similarly, 200,000 myosin 2A KO cells were transfected with 5 µg DNA. Four hours post-transfection, all cells were incubated overnight with 20 nM JFx650-HaloTag ligand (Lavis Lab, Lot: Sep-1-152).

### Generation of CRISPR knockin and knockout cell lines

Halo-2A knock-in cells were described previously (36). Briefly, immortalized parental fibroblasts (JR20) (37) were transfected with pSpCas9(BB)-2A-Puro (PX459) V2.0 (Plasmid 62988, Addgene) with target sequence 5’-AAACTTCATCAATAACCCGC-3’ generated using established protocols (38), and a donor plasmid (pUC57 with 794 bp 5’ HDR of genomic sequence immediate upstream of the endogenous start codon, a HaloTag with an 18 amino acid GS-rich linker, an 802 bp 3’ HDR of genomic sequence immediately downstream of the endogenous start codon with silent PAM site mutation). Single-cell sorting was performed 5–10 days after transfection. Individual clones were evaluated for knock-in via western blot analysis and microscopy. Clones used in this study include Halo-2A clone 2 (H2A2).

To generate myosin 2A KO lines, two lentiviral plasmids targeting Myh9 were constructed: pLenti-CRISPRvs-puro (Plasmid 98290, Addgene; target = ACCCTGCATCATGCTCCGGT-AGG) and pLenti-CRISPRvs-puro (Plasmid 98291, Addgene; target = GCAGCACCGAAGCTTCGTTG-AGG). Lentivirus was generated in HEK293 cells and placed directly on fibroblasts. Cells were selected in both puromycin and hygromycin and single cell sorting was performed to isolate knockout clones. Knockouts were validated using western blot analysis. JR20-myosin 2A-KO clone 3 was used for these experiments.

### Triton-fractionation assay

Cells were washed with PBS and imaged in cell media in 5% CO2 at 37°C. For Y27632 (688001, EMD Millipore) experiments, cells were treated with 10 µM of the drug for 30 minutes prior to imaging. Cells were then permeabilized in triton buffer (0.6% Triton X-100 (Millipore, 9002-93-1), 4% PEG 8000 (Promega, V3011), 5 mM NaCl, 140 mM potassium acetate, 100 mM PIPES, 1mM EGTA, 1 mM MgCl2) for 5 minutes before imaging the insoluble fraction.

### Western blotting

Cells were pelleted, resuspended in 2x Tris-Glycine Laemelli sample buffer (1610737, Biorad) supplemented with 10% beta-mercaptoethanol and 1x Halt protease & phosphatase inhibitor cocktail (78442, Thermo Scientific), and boiled for 5 minutes at 95°C. Cell lysates were analyzed with SDS–PAGE using 4-15% Mini-PROTEAN TGX Stain-Free Gels (4568083 Bio-Rad). Proteins were transferred to a nitrocellulose membrane (1704271, Biorad) using a Trans-Blot Turbo system (Biorad). Membranes were blocked for 1h at room temperature in 3% BSA/PBS supplemented with 0.1% Tween 20 (BP337-100, Fisher Scientific), incubated at 4°C overnight with primary antibody in 3% BSA/PBS + 0.1% Tween 20, washed three times for 5 min in blocking buffer, incubated for 1 h at room temperature with secondary antibody in blocking buffer, washed three times for 5 min in PBS + 0.1% Tween 20, rinsed in Milli-Q-H2O, and incubated with clarity Western ECL Substrate (1705061, Biorad) for 5 minutes and imaged on a ChemiDoc MP Imaging System (Biorad) for signal detection. The following primary antibodies were used: rabbit anti-beta actin (GTX109639, GeneTex, 1/5000 dilution), mouse anti-GFP (sc-9996, Santa Cruz, 1/2000 dilution) and rabbit anti-myosin IIA HC (MP3791, ECM Biosciences, 1/5000 dilution).

The following secondary antibodies were used at a 1/5,000 dilution: HRP goat anti-rabbit IgG (H+L) antibody (K1223, Apex bio), HRP goat anti-mouse IgG (H+L) antibody (K1221, Apex bio).

### Quantitative triton assays

Triton extraction assays were performed using a spinning disk confocal microscope from 3i (Intelligent Imaging Innovations) consisting of an Axio Observer 7 inverted microscope (Zeiss) attached to a W1 Confocal Spinning Disk (Yokogawa) with Mesa field flattening (Intelligent Imaging Innovations), a motorized X,Y stage (ASI), and a Prime 95B sCMOS (Photometrics) camera. Illumination was provided by a TTL triggered multifiber laser launch (Intelligent Imaging Innovations) consisting of six diode laser lines (405/445/488/514/561/640 nm) and all matching requisite filters. Temperature and humidity were maintained using a Bold Line full enclosure incubator (Oko Labs). The microscope was controlled using Slidebook 2023 Software (Intelligent Imaging Innovations). Cells were imaged using the 488 (EGFP constructs) and 640 (HaloTag) lasers at 50% laser power and 500 ms exposure time, using a 20 × 0.8 NA Plan-Apochromat objective (Zeiss).

All image analysis was performed in ImageJ-win64. Background images of cell media and triton buffer were taken on the spinning disk for both the 488 and 640 channels, using the same imaging setting described above. These images were darkfield corrected and the mean intensity was adjusted to 1. The raw data taken in cell media (pre-triton) and triton buffer (post-triton) were then divided by the cell media background and triton background images, respectively. All images were registered and tresholded to create a cell mask based on the 640 channel (Halo-2A). The masks were despeckled to reduce noise. All cells located at the border of the image, untransfected cells, or clustered cells were excluded from the analysis.

The resulting cell masks were dilated and the cell mask was subtracted from the dilated mask to create a donut mask. For each channel pre and post triton treatment, the median intensity within the donut mask (local background) was subtracted from the mean intensity within the cell mask. Finally, the insoluble fraction was calculated as the the post-extraction intensity/pre-extraction intensity ratio (Fig. 3 and 4). For the myosin 2A KO cells, the same image analysis pipeline was used to calculate the post triton-extraction intensity/pre triton-extraction intensity ratios., but the masks were based on the GFP signal.

### High Resolution Imaging

Halo-2A fibroblasts expressing EGFP-2A constructs were stained with JFx650-HaloTag ligand. Images were collected on a Zeiss LSM 880 Airsycan confocal in SR mode with a 63x 1.4 NA objective. Three slice z-stacks were collected at the ventral surface with optimal step size (0.17 µm). Images were processed in Zen Blue with “joint Deconvolution” processing using 10 iterations for EGFP channel and 20 iterations for JFx650-HaloTag channel. Maximum intensity orthogonal projections were generated using FIJI/ImageJ.

### Statistical analysis

Statistical analyses were performed using Prism (GraphPad). Data are presented as mean +/-standard deviation,and comparisons were made using values from each independent experiment. We performed nested one-way ANOVA and Tukey’s multiple comparison tests. P values <0.05 were considered significant. *, **, ***, **** indicate p ≤ 0.05, p ≤ 0.01, p ≤ 0.001, p ≤ 0.0001.

### Data availability

The data that support the findings of this study are available upon reasonable request from the corresponding author [Jordan Beach].

## Acknowledgements

Research reported in this publication was supported by the Maximizing Investigators’ Research Award (MIRA) (R35) from the National Institute of General Medical Sciences (NIGMS) of the National Institutes of Health (NIH) under grant number R35GM138183 to JRB. We thank Dr. Patrick Oakes, and the Beach and Oakes Labs for intellectual and emotional support throughout.

## Author contributions

JRB conceived the study. JRB and SS designed the experiments and analyses. KW, HP, HW, MAQ, MAB, SS and JRB performed the experiments and analyzed the data. KW, SS and JRB wrote the manuscript.

## Supplementary Tables

**Table S1.** Statistical comparisons using nested Anova or nested t-tests. Number of total cells and experimental replicates indicated in parenthesis. *, **, ***, **** indicate p ≤ 0.05, p ≤ 0.01, p ≤ 0.001, p ≤ 0.0001.

| EGFP constructs compared within experiment |  |  |  |
| --- | --- | --- | --- |
| (numbers indicate: nested anova;nested t-test) |  |  |  |
| Halo-2A Fibroblasts | EGFP-2A | EGFP-2A-ΔACD | EGFP-2A-ΔIQ2ΔACD |
| EGFP (n = 51, 3 replicates) | 0.0002***;0.0041** | 0.4647;<0.0001**** | 0.0002***;<0.0001**** |
| EGFP-2A (n = 46, 3 replicates) |  | 0.0008***;0.0098** | 0.9998;0.9531 |
| EGFP-2A-ΔACD (n = 39, 3 replicates) |  |  | 0.0009***;<0.0001**** |
| EGFP-2A-ΔIQ2ΔACD (n = 47, 3 replicates) |  |  |  |
| Myosin 2A KO | ΔACD | ΔIQ2ΔACD |  |
| EGFP-2A (n = 52, 4 replicates) | 0.0056**;0.009** | 0.0577;0.0547 |  |
| EGFP-2A-ΔACD (n = 23, 3 replicates) |  | 0.2591;0.0242* |  |
| EGFP-2A-ΔIQ2ΔACD (n = 27, 3 replicates) |  |  |  |
| ROCK inhibition | ΔACD | ΔIQ2ΔACD |  |
| EGFP-2A (n = 46, 4 replicates) | 0.5991;0.2621 | 0.5635;0.4058 |  |
| EGFP-2A-ΔACD (n = 31, 3 replicates) |  | 0.2001;0.1257 |  |
| EGFP-2A-ΔIQ2ΔACD (n = 28, 3 replicates) |  |  |  |
| Individual EGFP-2A constructs compared across experiments |  |  |  |
| EGFP-2A | Myosin 2A KO | ROCK inhibition |  |
| Halo-2A | 0.8379;0.5992 | 0.0246*;0.0217* |  |
| Myosin 2A KO |  | 0.0418*;0.0227* |  |
| EGFP-2A-ΔACD | Myosin 2A KO | ROCK inhibition |  |
| Halo-2A | 0.8861;0.6834 | 0.1157;0.0384* |  |
| Myosin 2A KO |  | 0.2547;0.2598 |  |
| EGFP-2A-ΔIQ2ΔACD | Myosin 2A KO | ROCK inhibition |  |
| Halo-2A | 0.0618;<0.0001**** | 0.0957;0.0973 |  |
| Myosin 2A KO |  | 0.9377;0.7919 |  |

## Supplementary Figures

**Fig. S1.**
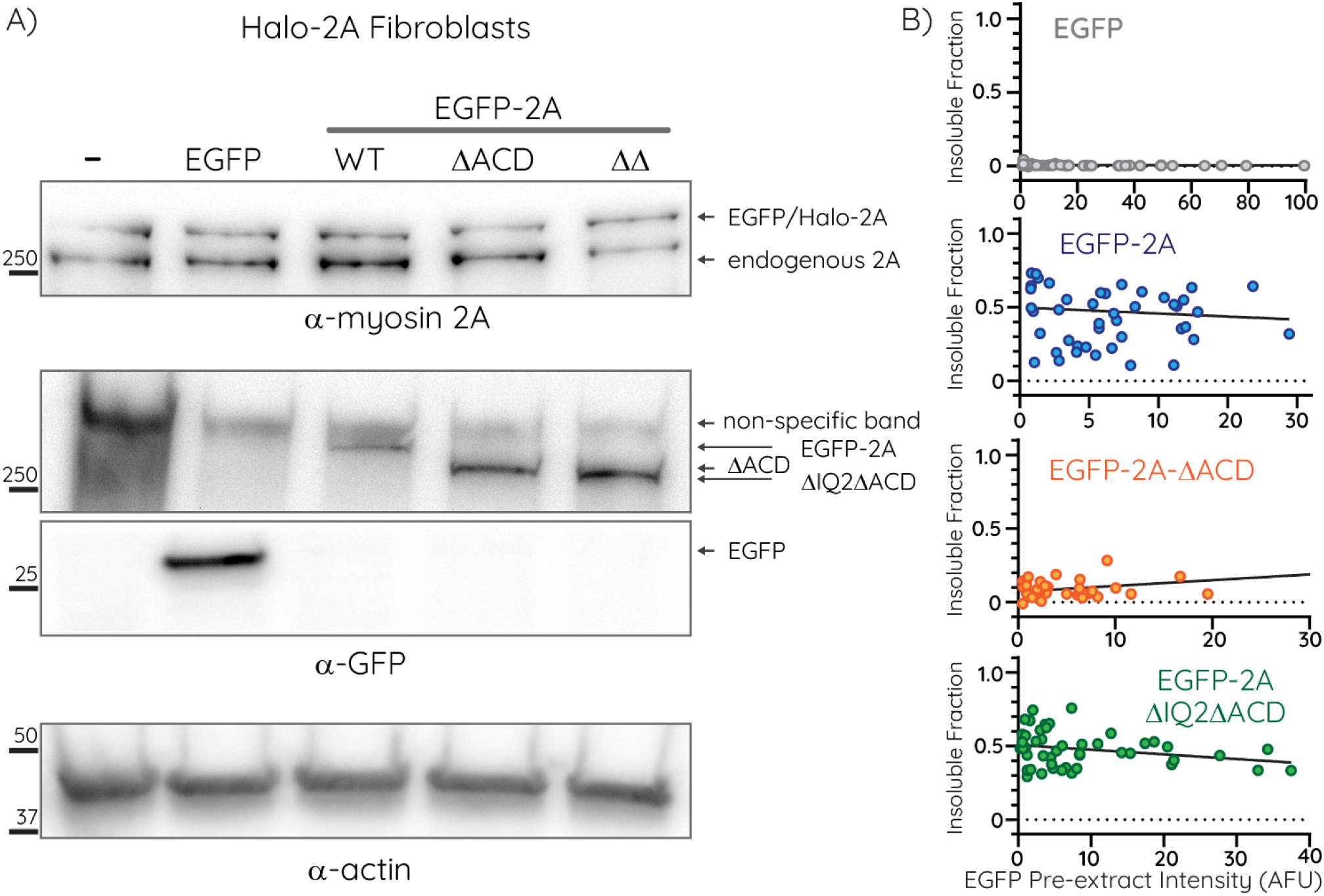
Expression analyses of exogenous EGFP constructs. A) Expression levels of the indicated EGFP-2A constructs in Halo-2A fibroblast lysates were determined via western blot analysis with the indicated antibodies. As exogenous EGFP-2A and endogenous Halo-2A have the same electrophoretic mobility, the extent of exogenous overexpression relative to endogenous cannot be determined. The EGFP-ΔACD truncation construct has a higher electrophoretic mobility (lower molecular weight) compared to full-length Halo-2A. However, the myosin 2A antibody was raised against a C-terminal epitope that is absent in the EGFP-2A-ΔACD constructs. B) Insoluble fractions from untreated Halo-2A fibroblasts in Fig. 3B plotted as a function of EGFP pre-extraction intensity. These data argue that the impact of variations in EGFP-2A overexpression levels on filament assembly is negligible.

**Fig. S2.**
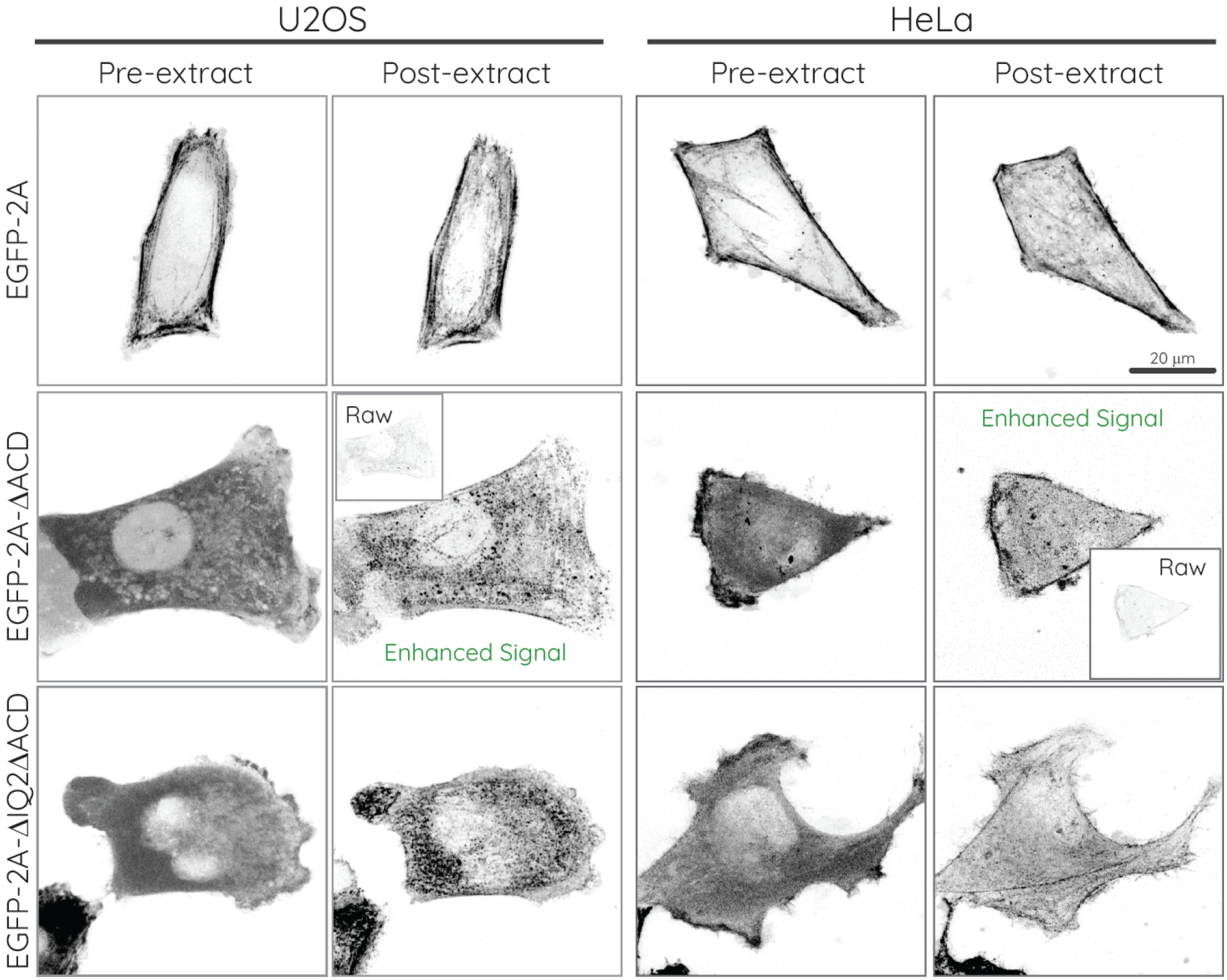
EGFP-2A-ΔACD and EGFP-2A-ΔIQ2ΔACD are also filamentous in U2OS and HeLa. Examples of summed confocal z-projections of U2OS or HeLa cells transiently overexpressing the indicated EGFP-2A constructs pre and post extraction. Extraction of EGFP-2A-ΔACD expressing cells diminished signal to near background (see Raw insets) but brightness enhancement revealed filamentous localization, consistent with these constructs co-assembling with endogenous myosin 2A regardless of cell type. Extraction of EGFP-2A and EGFP-2A-ΔIQ2ΔACD did not require brightness enhancement (pre- and post-extraction images scaled identically).

**Fig. S3.**
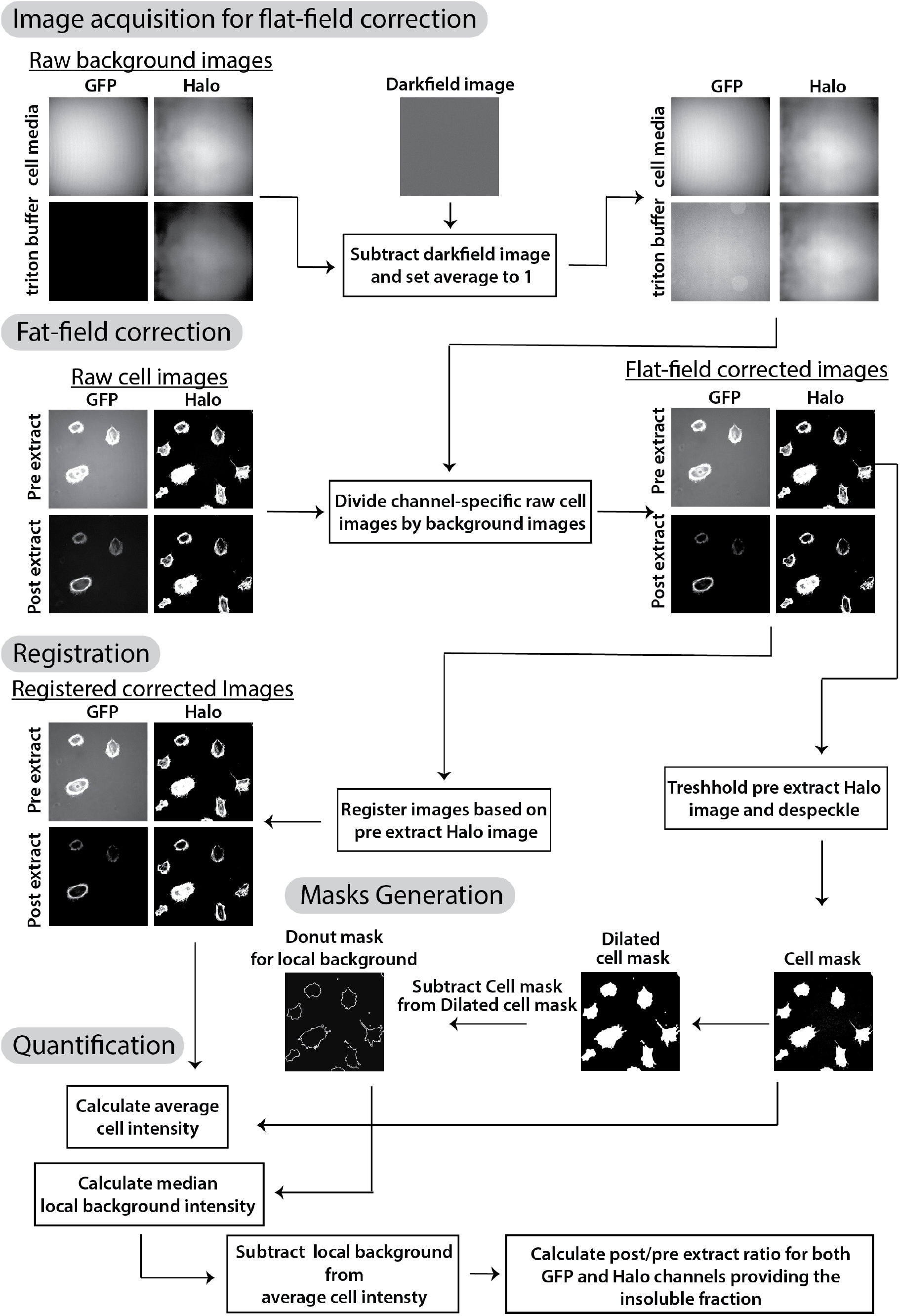
Analysis pipeline for imaging-based triton fractionation assay. See quantitative triton assay subsection in methods section for details.

## Supplementary Movie Legends

**Movie 01 - Time-lapse imaging EGFP constructs during triton extraction**. Fibroblasts expressing EGFP alone or EGFP-2A constructs as indicated were imaged on a Zeiss LSM 880 Airyscan confocal. A 3 *µ*m stack with 7 × 500 nm steps at the ventral surface was collected every 15 seconds and sum projected. LUT applied is in bottom right corner (MQ_div-magma). Triton extraction buffer was added after the first two time points. Video frame rate = 5 fps.

**Movie 02 - Time-lapse imaging of** Δ**IQ2**Δ**ACD**. A fibroblast expressing EGFP-2A-ΔIQ2ΔACD was imaged on a Zeiss LSM 880 Airyscan confocal. A 1.5 *µ*m stack with 4 × 500 nm steps at the ventral surface was collected every 30 seconds and sum projected. Magenta box on left indicates zoomed area on right. Video frame rate = 25 fps.

## Notes

### Competing Interest Statement

The authors have declared no competing interest.

